# KDM6A mediated expression of the long noncoding RNA DINO causes TP53 tumor suppressor stabilization in Human Papillomavirus type 16 E7 expressing cells

**DOI:** 10.1101/2020.01.01.892612

**Authors:** Surendra Sharma, Karl Munger

## Abstract

HPV16 E7 has long been noted to stabilize the TP53 tumor suppressor. However, the molecular mechanism of TP53 stabilization by HPV16 E7 has remained obscure and can occur independent of E2F regulated MDM2 inhibitor, p14^ARF^. Here, we report that the Damage Induced Noncoding (DINO) lncRNA (DINOL) is the missing link between HPV16 E7 and increased TP53 levels. DINO levels are decreased in cells where TP53 is inactivated, either by HPV16 E6, expression of a dominant negative TP53 minigene or by TP53 depletion. DINO levels are increased in HPV16 E7 expressing cells. HPV16 E7 causes increased DINO expression independent of RB1 degradation and E2F1 activation. Similar to the adjacent CDKN1A locus, DINO expression is regulated by the histone demethylase, KDM6A. DINO stabilizes TP53 in HPV16 E7 expressing cells and as a TP53 transcriptional target, DINO levels further increase. Similar to other oncogenes such as adenovirus E1A or MYC, HPV16 E7 expressing cells are sensitized to cell death under conditions of metabolic stress and in the case of E7, this has been linked to TP53 activation. Consistent with earlier studies, we show that HPV16 E7 expressing keratinocytes are highly sensitive to metabolic stress induced by the antidiabetic drug, metformin. Metformin sensitivity of HPV16 E7 expressing cells is rescued by DINO depletion. This work identifies DINO as a critical mediator TP53 stabilization and activation in HPV16 E7 expressing cells.

**IMPORTANCE:** Viral oncoproteins, including HPV16 E6 and E7 have been instrumental in elucidating the activities of cellular signaling networks including those governed by the TP53 tumor suppressor. Our study demonstrates that the long noncoding RNA DINO is the long sought missing link between HPV16 E7 and elevated TP53 levels. Importantly, the TP53 stabilizing DINO plays a critical role in the predisposition of HPV16 E7 expressing cells to cell death under metabolic stress conditions from metformin treatment.

## INTRODUCTION

The cancer-associated, high-risk human papillomavirus (HPV) E6 and E7 proteins have oncogenic activities and are necessary for tumor induction and maintenance. These low-molecular weight proteins lack intrinsic enzymatic activities and function by subverting host cellular regulatory networks (1, 2). Over the years, many potential cellular targets of the HPV E6 and E7 proteins have been identified. The main focus of these studies, however, has been on identifying host cellular protein targets and the resulting dysregulation of protein coding mRNAs. Protein coding genes, however, only constitute a minor fraction (<3%) of the human transcriptome (3), and advances in next generation sequencing technologies have shown that noncoding RNAs including microRNAs and long noncoding (lnc) RNAs, are also aberrantly expressed in HPV E6 and/or E7 expressing cells (4-7). It is now firmly established that noncoding RNAs are important regulators of a variety of cellular processes that drive specific biological phenotypes (8-11).

Many viral proteins, including the E6 and E7 proteins encoded by high-risk HPVs have evolved to modulate the TP53 tumor suppressor pathway (12). TP53 is critically involved in sensing many forms of cellular stress, including those caused by viral infection or nutrient starvation. Upon activation, TP53 is stabilized and elicits transcriptional programs that trigger cytostatic or cytotoxic responses, including cycle arrest, senescence or apoptosis (13). Hence, TP53 levels and activity are tightly regulated. A well-known regulator of TP53 is the MDM2 ubiquitin ligase. MDM2 is a TP53 responsive gene, which in turns binds and targets TP53 for ubiquitin dependent proteasomal degradation (14). MDM2 activity is regulated at multiple levels. One important inhibitor of MDM2 is the p14^ARF^ protein, which is regulated by E2F transcription factors, the downstream effectors of the retinoblastoma (RB1) tumor suppressor.

The E6 and E7 proteins encoded by high-risk HPVs, including HPV16, each modulate TP53 levels and activity. HPV16 E6 targets TP53 for proteasome mediated degradation through an MDM2 independent mechanism that involves the E6-associated ubiquitin ligase, UBE3A (E6AP) (15, 16). The ability of high-risk HPV E6 proteins to inhibit TP53 is thought to have evolved from the necessity to counteract TP53 stabilization and activation triggered by expression of the HPV E7 oncoprotein (17-22). Consequently, several other DNA tumor viruses, including adenoviruses and many polyomaviruses that encode proteins that inactivate the RB1 tumor suppressor also encode proteins that dampen TP53 tumor suppressor activity. In the case of the adenovirus E1A protein, TP53 stabilization was shown to be mediated by the E2F induced MDM2 inhibitor, p14^ARF^ (23). Surprisingly, however, HPV16 E7 has been shown to stabilize TP53 independently of p14^ARF^ (21) and, to this date, the mediator of HPV16 E7 mediated TP53 stabilization has remained elusive.

In recent years, it has been appreciated that various host cellular noncoding RNAs also function as important regulators of the TP53 tumor suppressor pathway. In addition to microRNAs (24), a variety of cellular lncRNAs have been identified as components of the TP53 signaling network (25). LncRNAs are defined as transcripts longer than 200 base pairs with no or limited (<100 amino acids) protein coding capacity. LncRNAs are versatile molecules and can form complexes with RNA, DNA and proteins. Some lncRNAs including lincRNA-p21, PURPL, MEG3 and DINO are transcriptional targets of TP53, and can also modulate TP53 levels and activity (26-29). DINO (Damage Induced Noncoding) lncRNA expression is induced by TP53 as a consequence of DNA damage. It has been reported that DINO binds TP53, thereby stabilizing and activating it. Hence, DINO functionally counteracts MDM2. In the present study, we report that DINO is expressed at high levels in HPV16 E7 expressing cells, and this upregulation correlates with the ability of HPV16 E7 to cause TP53 stabilization. High level DINO expression in HPV16 E7 expressing cells is driven by the KDM6A histone demethylase and TP53, and DINO, in turn, is necessary for maintaining elevated TP53 levels in HPV16 E7 expressing cells. Moreover, we show that the sensitivity of cells to undergo cell death in response to metabolic stress can be controlled by modulating DINO levels.

## RESULTS

### DINO levels correlate with TP53 levels in HPV oncoprotein expressing cells

Expression of high-risk HPV E6 and E7 proteins is known to modulate TP53 levels and activity. While HPV16 E7 expression causes TP53 stabilization, HPV16 E6 expression causes TP53 degradation. Hence, we determined whether the levels of DINO, a TP53 driven lncRNA correlate with TP53 levels in HPV16 oncogene expressing primary Human Foreskin Keratinocytes (HFKs). HFKs with expression of a dominant negative (dn)TP53 C-terminal fragment were used as a control (30). We assessed TP53 levels by immunoblotting and quantified DINO levels by quantitative reverse transcription polymerase chain reaction (qRT-PCR). Compared to control HFKs, we detected lower DINO levels in HFK populations where TP53 is inactivated either by the expression of the HPV16 E6 protein or by the dnTP53 C-terminal minigene (E6: 0.14 +/−0.01, p < 0.001, E6/E7: 0.22 +/−0.04, p < 0.001 and dnTP53: 0.28 +/−0.01, p < 0.001) (Figure 1B). This is consistent with a previous study that showed that DINO expression is controlled by TP53 (28). In contrast, HPV16 E7 expressing HFKs, which, as published previously (17, 20, 21), express high levels of TP53 (Figure 1A), expressed significantly increased DINO levels (E7: 6.14 +/−0.14, p < 0.001) (Figure 1B). Hence, DINO levels correlate with TP53 levels in HPV oncoprotein expressing cells.

**Figure 1:**
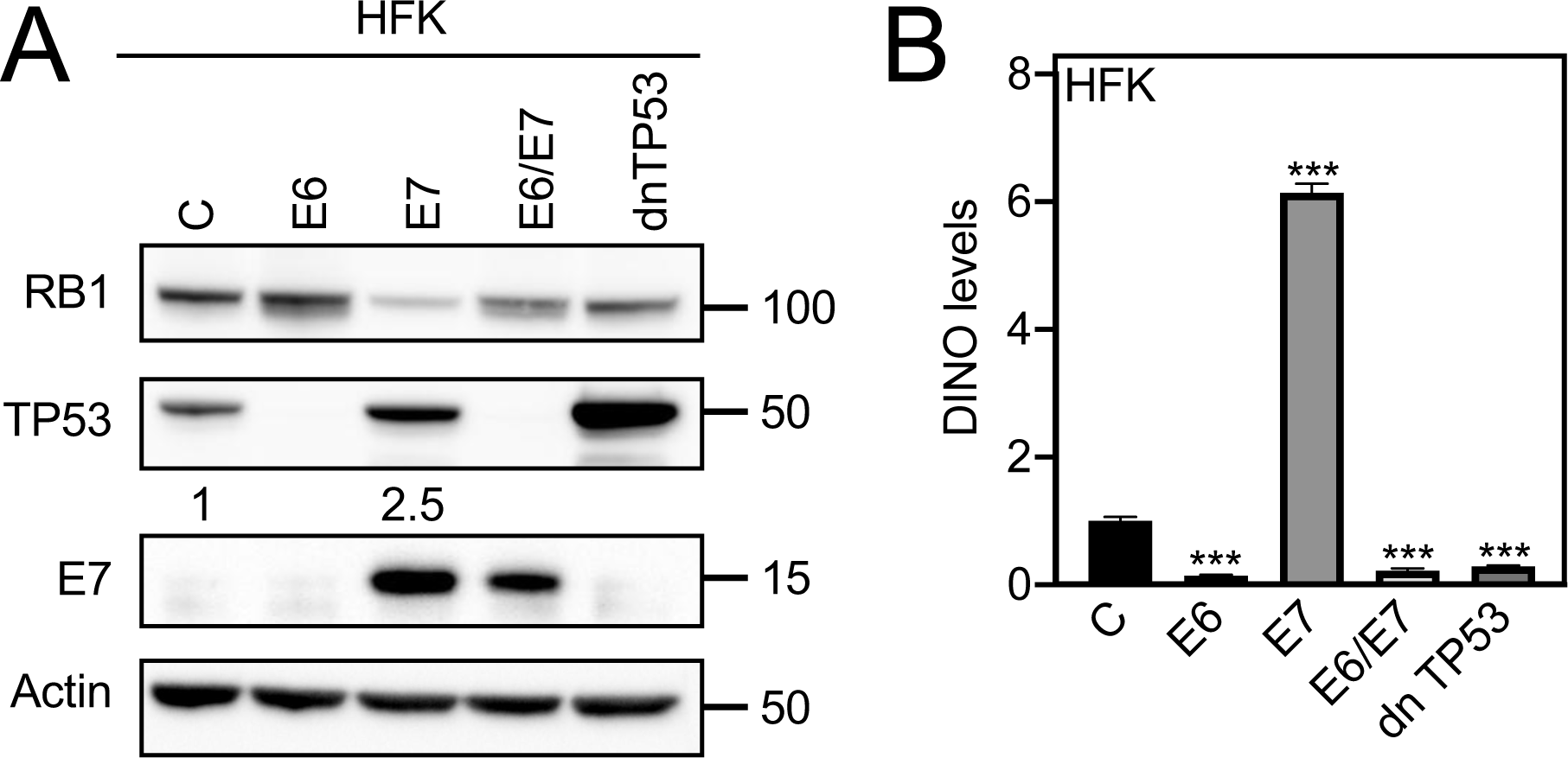
DINO levels corelate with TP53 levels in HPV oncoprotein expressing cells. HFK populations with stable expression of HPV16 E6 and/or E7 or a dominant negative TP53 minigene (dn TP53) were validated by determining the levels of RB1, TP53 and E7 by western blotting (A). DINO levels were accessed in these HFK populations by qRT-PCR assays (B). Levels shown are relative to control vector transduced HFKs. The bar graph depicts means ± SEM (n=3) calculated from a single representative experiment. *** p < 0.001 (Student’s t-test). Similar results were obtained in three independently derived HFK populations.

### TP53 driven DINO causes elevated TP53 levels in HPV16 E7 expressing cells

Given the correlation between DINO and TP53 levels in E7 expression cells, we next determined whether DINO expression in HPV16 E7 expressing cells was driven by TP53. We transiently depleted TP53 in control as well as in HPV16 E7 expressing human telomerase immortalized HFKs (iHFKs) with a pool of TP53 targeting siRNAs. To assess the effect of TP53 depletion on expression of TP53 target genes, we analyzed expression of CDKN1A (p21^CIP1^) by western blotting and qRT-PCR. As expected, CDKN1A protein and mRNA levels were decreased in control and HPV16 E7 expressing cells upon TP53 depletion (Figure 2A, B). Similarly, TP53 depletion also caused a significant decrease of DINO in control as well as in HPV16 E7 expressing cells (C: 0.09 +/−0.01, p < 0.001 and E7: 0.08 +/−0.02, p < 0.001) (Figure 2B).

**Figure 2:**
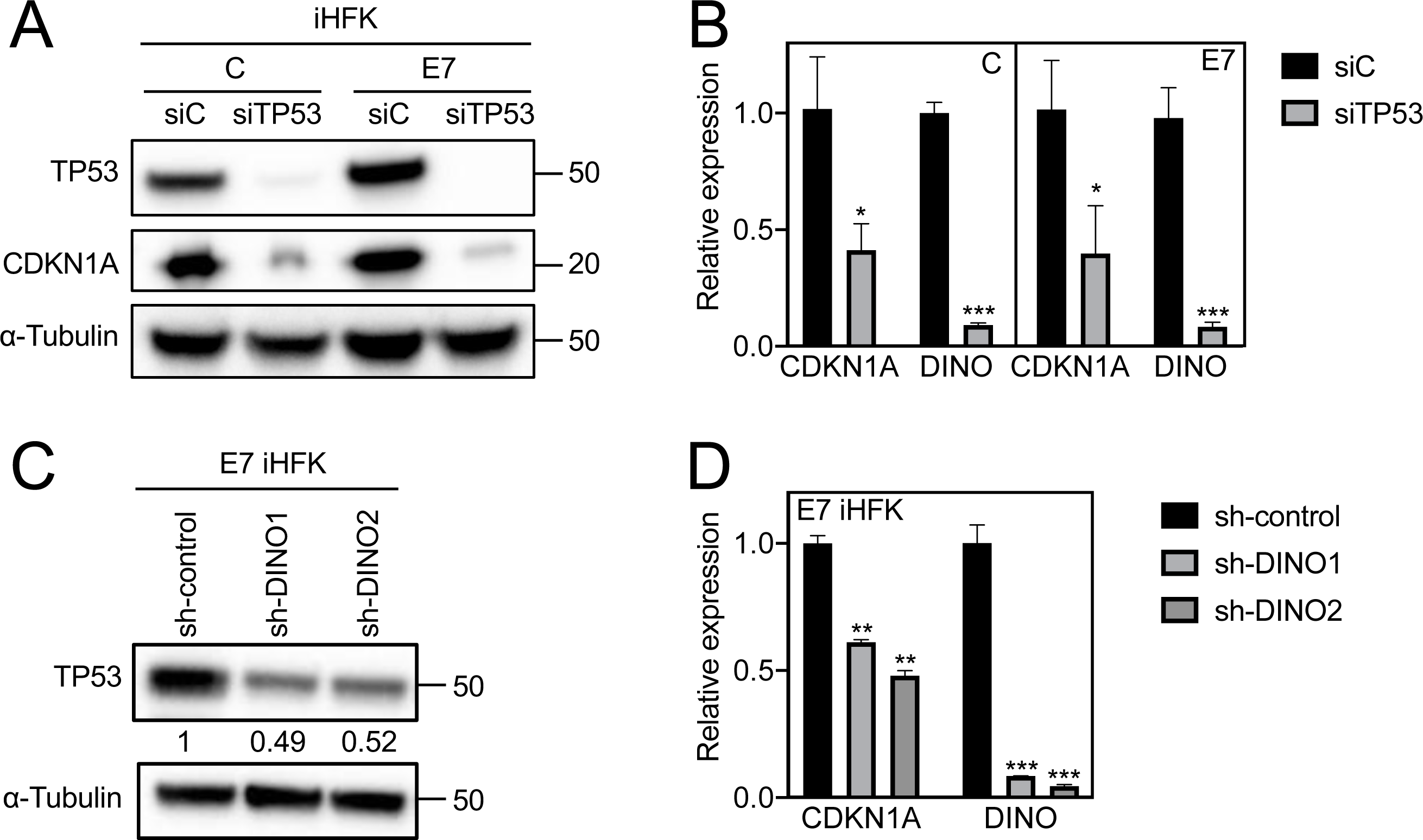
DINO causes elevated TP53 levels in HPV16 E7 expressing cells. TP53 depletion in both control (C) and HPV16 E7 (E7) expressing iHFKs were validated by western blotting and assessment of the canonical TP53 transcriptional target CKDN1A protein (A) and mRNA levels (B). DINO levels were determined in TP53 depleted iHFKs by qRT-PCR assays and results are shown by comparing it with control siRNA transfected iHFKs. For acute depletion of DINO, HPV16 E7 expressing iHFKs harboring doxycycline inducible DINO shRNA expression vectors were treated with 1μg/mL doxycycline for 48 hours. HPV16 E7 iHFKs with acute expression of non-targeting shRNA sequences were used as negative controls. Levels of TP53 protein were determined by western blotting (C). Expression of the TP53 responsive CDKN1A mRNA and DINO depletion were assessed by qRT-PCR (D). Bar graphs depict means ± SEM (n=3) calculated from a single representative experiment. *** p < 0.001, ** p < 0.01, * p < 0.05 (Student’s t-test). Similar results were obtained in three independent experiments.

To determine whether DINO may be the “missing link” that mediates TP53 stabilization in E7 expressing cells, we acutely depleted DINO in HPV16 E7 expressing iHFKs using doxycycline regulated shRNA expression. Depletion of DINO after doxycycline treatment for 48 hours by two different shRNAs each caused an approximately 50% decrease in TP53 levels in E7 expressing cells (Figure 2C). Similarly, mRNA levels of the TP53 transcriptional target gene, CDKN1A gene were also decreased (sh-DINO1: 0.61 +/−0.01, p < 0.01, and sh-DINO2: 0.48 +/−0.02, p < 0.01) (Figure 2D). Hence, DINO levels accumulate in HPV16 E7 expressing cells as a result of TP53 activation, and, in addition, they amplify TP53 stabilization and presumably TP53 activation in HPV16 E7 expressing cells.

### TP53 and KDM6A, but not RB1 and E2F1 are the upstream regulators of DINO

Previous mutational studies with HPV16 E7 revealed that TP53 stabilization by HPV16 E7 required similar sequences as those that are necessary for RB1 destabilization (20). Consistent with these earlier studies, the RB1 binding and degradation defective HPV16 E7 ∆DLYC mutant failed to stabilize TP53 (Figure 3A). Similarly, unlike wild type HPV16 E7, expression of HPV16 E7 ∆DLYC also failed to upregulate DINO in IMR90s normal human diploid lung fibroblasts (E7: 7.03 +/−1.28, p < 0.01 and ∆DLYC: 1.00 +/−0.18, non-significant) (Figure 3B). To determine whether DINO levels in HPV16 E7 expressing cells were increased as a consequence of RB1 tumor suppressor degradation and E2F activation, we transiently depleted RB1 and E2F1 in control and HPV16 E7 expressing iHFKs by transfecting the corresponding siRNA pools. Transfections of a TP53 targeting siRNA pool and a non-targeting siRNA pool were used as positive and negative controls, respectively. Depletion of the corresponding proteins was assessed by western blotting (Figure 3C). Whereas, as expected, TP53 depletion caused a significant decrease of DINO levels, RB1 and E2F1 depletion did not cause significant changes of DINO levels in control (siC: 1.00 +/−0.12; siRB1: 1.15 +/−0.03, p = non-significant (ns) and siE2F1: 0.96 +/−0.14, p = ns) or HPV16 E7 expressing cells (siC: 6.64 +/−0.52, siRB1: 5.58 +/−0.56, p = ns and siE2F1: 6.15 +/−0.04, p = ns) (Figure 3D). The DINO locus is in close proximity of the CDKN1A locus (28), which is subject to epigenetic de-repression by the histone H3 lysine 27 (H3K27) demethylase, KDM6A (31, 32). Hence, we next tested whether DINO expression is also regulated by KDM6A. Consistent with previous reports (32, 33), we found that KDM6A expression was increased in E7 expressing cells. Moreover, DINO expression was significantly decreased upon KDM6A depletion (C: 0.61 +/−0.09, p < 0.01 and E7: 2.83 +/−0.17, p < 0.001) (Figure 3D). Hence, HPV16 E7 causes increased DINO expression through a mechanism that involves KDM6A mediated de-repression.

**Figure 3:**
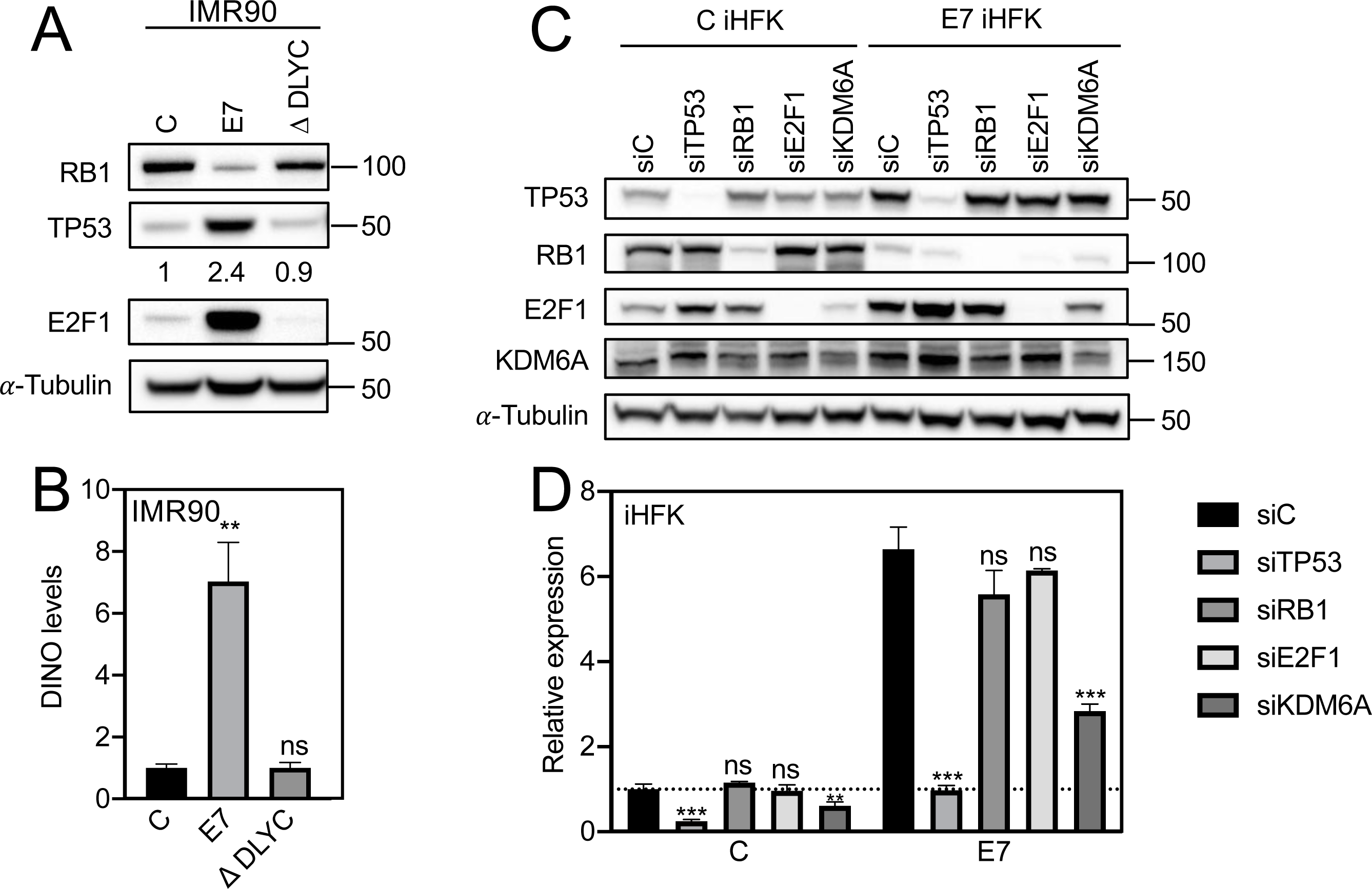
KDM6A and TP53, but not RB1 and E2F1 are the upstream regulators of DINO. IMR90 cell populations expressing either control, wild type HPV16 E7 or the RB1 binding/degradation defective HPV16 E7 ∆DLYC mutant, were assessed for RB1 degradation, TP53 stabilization and E2F1 expression by western blotting (A). DINO levels were accessed in these IMR90s populations by qRT-PCR (B). Levels shown are relative to control vector transduced IMR90s. Validation of TP53, RB1, E2F1 and KDM6A depletion in control (C) and HPV16 E7 (E7) expressing iHFKs by western blotting (C). DINO levels in each of these lines as determined by qRT-PCR (D). Bar graphs represent means ± SEM (n=3) calculated from a single representative experiment are shown. *** p < 0.001, ** p < 0.01 and ns = non-significant (Student’s t-test). Similar results were obtained in three independent experiments.

### Heightened metformin sensitivity of HPV16 E7 expressing cells is reversed upon depletion of DINO

Our research group has previously reported that HPV16 E7 expressing fibroblasts are sensitized to undergo cell death under conditions of serum starvation. This response was shown to be TP53 dependent since it was abrogated by co-expression of the HPV16 E6 protein or a dominant negative TP53 minigene (18, 20). Given our results that DINO is critical to TP53 stabilization and activation in HPV16 E7 expressing cells, we next wanted to determine whether vulnerability of HPV16 E7 expressing cells could be controlled by modulating DINO levels. Since keratinocytes are grown in serum free media, we first needed to define conditions that may mimic growth factor deprivation in keratinocytes. We evaluated metformin, a drug that induces metabolic stress by activating AMPK, inhibiting mTOR and mitochondrial respiratory complex I, and depleting glycolytic and TCA cycle intermediates (34-37). Our results show that HPV16 E7 expressing HFKs are more vulnerable than control HFKs when subjected to metformin treatment (C: 24.78 +/−1.12 and E7: 68.48 +/−0.38, p < 0.001) (Figure 4A). Heightened metformin vulnerability of HPV16 E7 expressing cells was abrogated upon HPV16 E6 co-expression (−0.46 +/−0.20, p < 0.001), consistent with the model that the vulnerability to metformin may be a consequence of E7 mediated TP53 stabilization and activation. To determine whether DINO may modulate metformin induced cell death, we depleted DINO in HPV16 E7 expressing primary HFKs and treated them with metformin. As expected, DINO depletion caused reduced levels of TP53 and decreased expression of the TP53 transcriptional target CDKN1A (Figure 4C, D). DINO depletion did not affect the viability of HPV16 E7 expressing HFKs when grown under standard tissue culture conditions. In contrast, however, DINO depleted HPV16 E7 expressing HFKs were significantly more resistant (sh-control: 50.76 +/−0.78, sh-DINO1: −1.71 +/−0.47, p < 0.001 and sh-DINO2: −10.21 +/−0.68, p < 0.001) to metformin treatment than normal HPV16 E7 expressing HFKs (Figure 4B). Hence, metformin can be utilized to investigate the metabolic vulnerability of HPV16 E7 expressing cells and DINO regulates cellular response to metformin induced metabolic stress.

**Figure 4:**
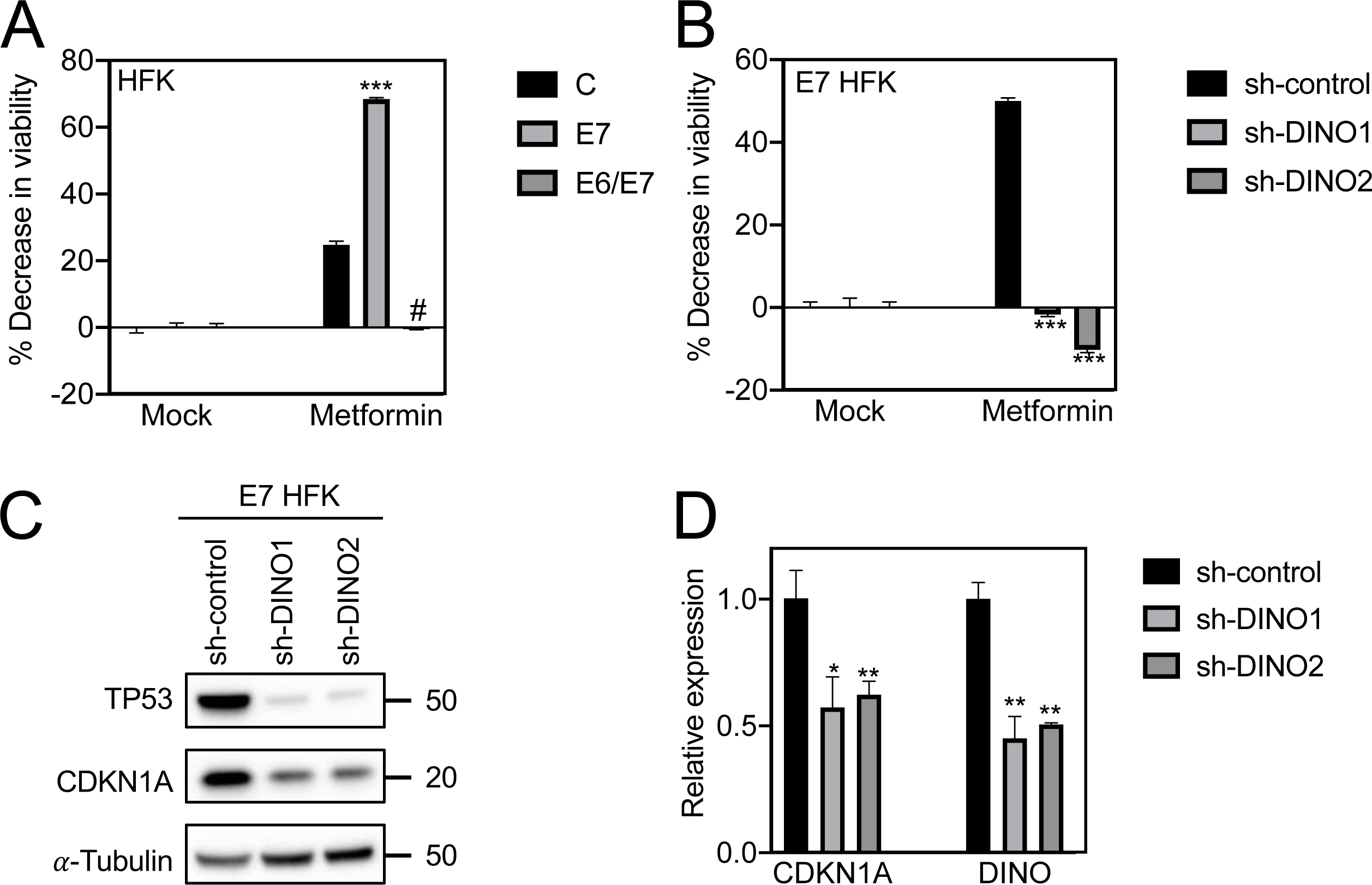
Heightened metformin sensitivity of HPV16 E7 expressing cells is reversed upon DINO depletion. Cell viability as assessed by the resazurin assay of control, HPV16 E7 and HPV16 E6/E7 expressing HFKs that were either untreated (mock) or treated with 20 mM metformin for 96 hours (A). Viability of HPV16 E7 expressing HFKs with expression of either of two DINO specific shRNAs or a scrambled control shRNA (B). TP53 and CDKN1A expression in these cell lines as determined by western blotting (C). Expression of CDKN1A mRNA and validation of DINO depletion in these HFK populations as determined by qRT-PCR (D). Bar graphs depict means ± SEM (n=3) calculated from a single representative experiment. *** p < 0.001, ** p < 0.01, * p < 0.05, # p < 0.001 (Student’s t-test). Similar results were obtained in three independently derived HFK populations.

## DISCUSSION

The TP53 tumor suppressor is a central hub that integrates various cellular stress signals and triggers appropriate cytostatic and cytotoxic responses. In normal cells, TP53 has a very short half-life and is present at low levels because the TP53 regulated MDM2 ubiquitin ligase targets TP53 for proteasomal degradation. Oncogenic insults can trigger TP53 activation. In response to RB1 tumor suppressor inactivation, the E2F regulated p14^ARF^ protein inhibits MDM2 and causes TP53 activation. However, it became clear early on that there must be additional regulators of TP53, since p19^ARF^ null mouse embryo fibroblasts were shown to still be capable of TP53 stabilization and activation in response to DNA damage (38). Similarly, even though HPV16 E7 causes E2F activation, it can stabilize and activate TP53 in p19^ARF^ null mouse embryo fibroblasts and the mechanism of HPV16 E7-mediated TP53 stabilization and activation has remained elusive (21).

DINO is a TP53 responsive gene and DINO levels increase in response to DNA damage. DINO was reported to bind and stabilize TP53, thereby amplifying the TP53 transcriptional response to DNA damage. Hence, DINO functionally counteracts MDM2 (28, 39). Here, we show that DINO expression correlates with TP53 expression in HPV16 oncoprotein expressing cells and is highly expressed in HPV16 E7 expressing keratinocytes. DINO depletion causes a decrease in TP53 levels in HPV16 E7 expressing cells and, thus, DINO is the missing link between HPV16 E7 expression and TP53 stabilization. The RB1 binding/degradation defective HPV16 E7 ∆DLYC mutant, which was previously shown to be defective for TP53 stabilization (20), does not cause increased DINO expression. Hence, we were quite surprised that we did not detect any changes in DINO expression when RB1 was depleted by RNAi. This suggests that RB1 degradation and the ensuing activation of E2F are not the primary triggers of DINO expression. In contrast, however, depletion of the H3K27 demethylase KDM6A, which is expressed at higher levels in E7 expressing cells, causes a significant decrease in DINO. Hence our results, suggest a model whereby E7 mediated upregulation of KDM6A provides the initial trigger for DINO induction. Our current results cannot distinguish, however, whether, similar to the proximal CDKN1A locus (28, 32), DINO expression is directly de-repressed through KDM6A mediated H3K27 demethylation, or whether this involves an indirect mechanism. It is noted, however, that DINO and CDKN1A are distinct genetic loci and are expressed from opposite strands in divergent directions (28). According to our model, the initial KDM6A mediated increase in DINO expression causes TP53 stabilization and activation, which in turn causes a further increase in DINO and TP53. Since E7 will also cause increased expression of p14^ARF^ as a consequence of RB1 degradation and E2F activation, this may impair MDM2 activity and cause a further increase in TP53 and DINO (Figure 5). Importantly, this model, where the initial trigger of DINO expression is through KDM6A, accommodates the finding E7 expression can cause increased TP53 levels in p19^ARF^ deficient mouse embryo fibroblasts (21).

**Figure 5:**
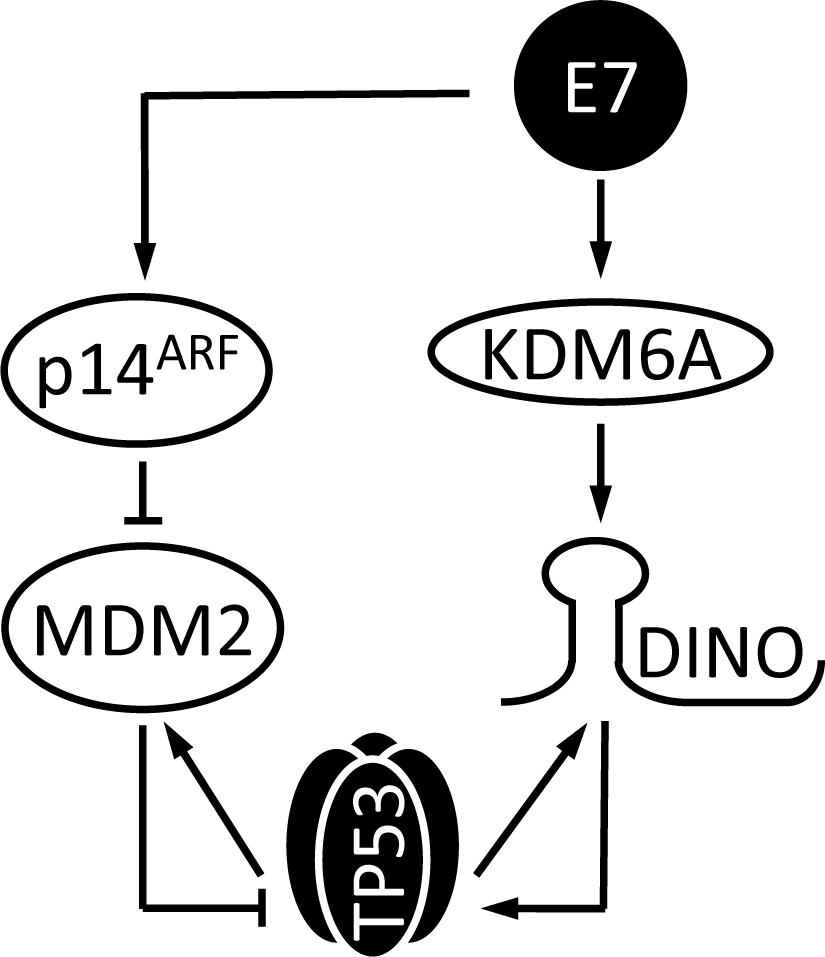
Model of TP53 stabilization and activation by the HPV16 E7 oncoprotein. HPV16 E7 expression triggers DINO expression through a KDM6A mediated pathway. This causes TP53 activation which results in further TP53 mediated DINO transcription. HPV16 E7 also induces p14^ARF^ as a consequence of RB1 degradation which inhibits MDM2 ubiquitin ligase activity causing a further increase in TP53 and DINO. See text for details.

Having established DINO as the key mediator of elevated TP53 levels in E7 expressing cells, we next wanted to determine whether DINO may not only modulate TP53 levels but also a known TP53 dependent biological phenotype of HPV16 E7 expressing cells. We previously discovered that, similar to adenovirus E1A or MYC, HPV16 E7 expressing primary cells undergo cell death when deprived of growth factors (17, 20, 40-42). This form of cell death is triggered by conflicting growth signals; the proliferative signal generated by oncogene expression that clashes with the antiproliferative signal as a consequence of serum deprivation. This likely represents a cell-intrinsic, innate tumor suppressive response that has been dubbed the “trophic sentinel response” (43). In the case of HPV16 E7, this response is TP53 dependent and is overridden by co-expression of the HPV16 E6 oncoprotein (18, 20).

Serum deprivation is not a practicable approach to induce metabolic stress in keratinocytes, the normal host cells of HPVs, since they are grown in serum free media. Hence, we evaluated other means of generating metabolic stress and used metformin, a member of anti-diabetic biguanide compounds that are evaluated for repurposing as cancer therapeutic and/or chemoprevention drugs. Metformin treatment of control and HPV16 E7 expressing keratinocytes revealed that, compared to parental cells, HPV16 E7 expressing cells are more sensitive to metformin treatment and that DINO depletion abrogates this sensitivity.

Similar to microRNAs, lncRNA activity can be modulated by nucleic acid based inhibitors and mimics and it will be interesting to determine whether DINO based compounds may be developed that could be used to modulate the clinical response to drugs that cause metabolic stress in cancer cells, particularly in cancer such as HPV associated cancer where TP53 is not mutated.

## MATERIALS AND METHODS

### Cell culture

Primary human foreskin keratinocytes (HFKs) were prepared from a pool of 3 to 5 neonatal foreskins obtained anonymously from the Ob/Gyn department at Tufts Medical Center. Human telomerase (hTERT)-immortalized HFKs (iHFKs) were a generous gift from Dr. Aloysius Klingelhutz (44). HFKs and iHFKs were grown and maintained in keratinocyte-serum-free media (KSFM) supplemented with human recombinant epidermal growth factor and bovine pituitary extract (Invitrogen). IMR-90s normal human diploid lung fibroblasts were purchased from ATCC and maintained in Dulbecco’s Modified Eagle Medium (DMEM) supplemented with 10% fetal bovine serum. HPV 16 E6 and/or E7 and dominant negative (dn)TP53 C-terminal minigene expressing cells were created by retroviral infection with recombinant LXSN based retroviruses provided kindly by Dr. Denise Galloway (HPV oncogenes (45, 46)) and Dr. Moshe Oren (dnTP53 (30)). Cells were selected in 300 μg/ml G418 (Gibco) and G418-resistant pooled cell populations were used for analysis. In all HFKs studies, donor and passage matched HFKs of passages less than 8 times were used. In IMR90s studies, passage matched IMR90s of passages less than 10 times were used. Doxycycline and polybrene were purchased from Sigma. Metformin was purchased from Cayman Chemicals. Doxycycline and metformin were dissolved in PBS and freshly prepared for each use.

### Western blotting and antibodies

Lysates were prepared by incubating the cells in RIPA lysis buffer supplemented with Pierce protease inhibitor (Thermo Scientific) and Pierce phosphatase inhibitor (Thermo Scientific) at 4 C for 30 min. Proteins extracts were cleared by centrifugation at 15,000 g for 15 min. Equal amounts (50 μg) of proteins were loaded and fractionated on 4–12% NuPAGE Bis-Tris Gels (Invitrogen). Protein were transferred to PVDF membranes (Millipore) and blocked with TNET buffer (200 mM Tris–HCl, 1 M NaCl, 50 mM EDTA, with 0.1% Tween-20, pH 7.5) containing 5% nonfat dry milk at room temperature for 1 hour. Blots were incubated with following primary antibodies at 4 C overnight: TP53 (OP43, Calbiochem, 1:1000), RB1 (Ab-5, Millipore, 1:100), E2F1 (sc-251, Santa Cruz, 1:500), KDM6A (ab36938, Abcam, 1:300), CDKN1A (ab109520, Abcam, 1:1,000), α-tubulin (ab18251, Abcam, 1:1000) and actin (Ab-1501, Millipore, 1:1000). Membranes were rinsed in TNET and incubated with the corresponding secondary antibodies, horseradish peroxidase (HRP)-conjugated anti-mouse antibody (NA931, GE Healthcare Life Sciences, 1:10,000) or HRP-conjugated anti-rabbit antibody (NA934V, GE Healthcare Life Sciences, 1:10,000) for one hour at room temperature. Antigen/antibody complexes were visualized by Enhanced Chemiluminescence (Life Technologies) and signals were digitally acquired on a G:Box Chemi-XX6 imager with GeneSys software (Syngene). Protein bands were quantified using GeneTools Software (Syngene).

### Lentiviral expression plasmids

Lentiviral vectors encoding two separate DINO targeting shRNA hairpin sequences were kind gifts from Dr. Howard Chang (28). Doxycycline-inducible DINO knockdown vectors were created by inserting these DINO targeting hairpin sequences into the Tet-pLKO-puro vector backbone (Addgene plasmid #21915, (47)). Oligonucleotide sequences used for cloning were, sh-DINO1 oligo (5’-3’): ccggcacagaagaattggacattgaactcgagttcaatgtccaattcttctgtgttttt, and sh-DINO2 oligo (5’-3’): ccggctggtttatggagatgacataactcgagttatgtcatctccataaaccagttttt. As a negative control, non-mammalian shRNA sequences (Sigma SHC002) were cloned into Tet-pLKO-puro vectors by using oligo (5’-3’): ccggcaacaagatgaagagcaccaactcgagttggtgctcttcatcttgttgttttt. All inserted sequences were verified by DNA sequencing.

### RNA interference and lentiviral transduction

3×10^6^ control and HPV16 E7 iHFKs were seeded on 10 cm cell culture dish and allowed to adhere overnight. Next day, cells were transfected with 30 nM siRNAs specific for targeted genes (ON-TARGETplus SMARTpools; Thermo Scientific Dharmacon) or a negative control siRNA (Non-Targeting Pool; Thermo Scientific Dharmacon) using Lipofectamine RNAiMax reagent (Invitrogen) per the manufacturer’s instructions. Dharmacon references for gene specific siRNAs used in this study are, TP53 (L-003329-00), RB1 (L-003296-02), E2F1 (L-003259-00), KDM6A (L-014140-01) and non-targeting control (D-001810–10). Recombinant lentiviruses expressing either non-targeting control or DINO shRNA sequences were made by transfecting HEK293T cells with the corresponding lentiviral vector, psPAX2 packaging (Addgene#12260) and pMD2.G (Addgene#12259) envelope plasmid DNA at a ratio of 4:3:1, respectively. Culture medium was collected at 48 hours post transfection and used for infection in conjunction with 0.4 ug/mL polybrene. Post 4 hours of infections, the inoculum was removed and replaced with fresh media. Stable cell populations were generated post selection in 1μg/ml puromycin.

### RNA isolation and quantitative PCR

Total RNA was isolated using the Quick-RNA MiniPrep (Zymo Research) and 1 μg of total RNA was reverse transcribed using the Quantitect Reverse Transcription Kit (Qiagen) per the manufacturer’s instructions. Quantitative PCR (qPCR) was performed in triplicate using SYBR Green PCR Master Mix (Applied Biosystems) reagents in a StepOne Plus (Applied Biosystems) thermocycler system. For all qPCR reactions in this study, thermocycler settings of 20 sec at 95°C, followed by 40 cycles of 3 sec at 95°C and 30 sec at 60°C were used. The following qPCR primer sequences were used in this study. DINO: 5’-ggaggcaaaagtcctgtgtt −3’(forward) and 5’-gggctcagagaagtctggtg −3’(reverse); CDKN1A: 5’-catgtggacctgtcactgtcttgta −3’(forward) and 5’-gaagatcagccggcgtttg −3’(reverse); RPLP0:5’-atcaacgggtacaaacgagtc −3’(forward) and 5’-cagatggatcagccaagaagg −3’(reverse). The expression data shown was quantified using the ΔΔCT method by normalizing all the qPCR targets against a housekeeping gene, RPLP0.

### Cell viability

Cell viability was assessed using a resazurin assay, as previously described (48). At the day of cell viability reading, cells were incubated with fresh media containing 10 μg/ml resazurin (Sigma). After a one-hour incubation, fluorescence readings were recorded on a Synergy H1 microplate reader (BioTek) using 560 nm excitation and 590 nm emission filters.

## ACKNOWLEDGMENTS

We thank Dr. Howard Chang (Stanford University) for generously sharing DINO reagents, Dr. Al Klingelhutz (University of Iowa) for providing telomerase immortalized human foreskin keratinocytes and Drs. Amy Yee, Philip Hinds, Peter Juo, Claire Moore and members of the Munger Lab for stimulating discussions and valuable suggestions throughout the course of this work. Supported by PHS grants AI147176, CA066980 and CA228543 (K.M.) and a Dean’s Fellowship and a Tufts Collaborative Cancer Biology Award from the Tufts Graduate School of Biomedical Sciences (S.S.).

